# Playing a P300-based BCI VR game leads to changes in cognitive functions of healthy adults

**DOI:** 10.1101/2020.05.28.118281

**Authors:** Matvey Bulat, Alexandra Karpman, Alina Samokhina, Alexander Panov

## Abstract

In this paper, we present the results of a study to determine the effect of the P300-based brain-computer interface (BCI) virtual reality game on the cognitive functions of healthy human subjects. This study is a part of on-going research related to evaluation of the the long-term effect of P300 training in Virtual Reality surrounding (VR game) on the cognitive performance of the young healthy population. A comparison of results between 3 groups of participants (15 people each) revealed the progressing difference in cognitive assessment for experimental group played P300 BCI VR game, showing the positive increase in flanker and conjunction visual search task performance associated with selective attention and mental inhibition. We show that the effect is due to the use of P300 BCI paradigm. Our results suggest that P300 BCI games combined with virtual reality can not only be used for rehabilitation in patients with slight mental disorders or elderly, but for increasing some cognitive functions in healthy subjects, giving an additional improvement in learning in case of combination with possible educational tasks or used for attention training

**GRAPHICAL ABSTRACT:** Please check the journal’s author guildines for whether a graphical abstract, key points, new findings, or other items are required for display in the Table of Contents.

## 1 INTRODUCTION

### 1.1 BCI

A brain-computer interface is a communication system that provides the user with the ability to send signals (commands or messages) to external world directly from the brain, by using only the electric activity of brain cells populations or single cells (Wolpaw et al., 2002). Research in the area of brain-computer interfaces is ongoing both in academia and industry. In the recent years many neurotechnological companies were founded (e.g Neuralink, Kernel, Paradromics, BrainCo, etc.), or got internal neurotech daparmtments built from startups (e.g. Ctrl Labs at Facebook).

While a lot of research of academic groups is focused on the medical field, investigating possible ways of diagnostics of neurological disorders, treatment procedures (Bockbrader et al., 2018) (in combination with simultaneous stimulation techniques: TMS, TdCS, etc.), and rehabilitation protocols for patients, affected by stroke, spinal cord injury, paralysis, plegia etc.; companies are more dedicated to the development of a more user-friendly and convenient to use BCIs which can allow to monitor cognitive functions, mental control, attention levels etc.

Close interaction between research groups and business is nowadays becoming a keystone in productive boosting in fundamental research, hardware and software evolution and conscious development of the whole neurotech field, leading to increase in human well being levels and to understanding the aspects of brain functionality at neuro-physiological level.

### 1.2 Games

For consumer-affordable BCIs, electroencephalography (EEG) is the most suitable technique, allowing non-invasive brain signal acquisition, however some other modalities such as near-infrared spectroscopy (NIRS) or magnetoencephalography (MEG) are also promising to become consumer-available in the next years (Yang et al., 2019; Boto et al. 2018). EEG measures electrical activity from the brain via multiple electrodes placed across the scalp. It is low-cost and has high temporal resolution (few ms), which allows one to get and process the signal in real-time, allowing to build adaptive BCI.

Aside from medical applications an intriguing sphere for EEG-based BCI implementation is gaming. Game industry is able not only to gain from BCIs adaptive gameplay and new ways of interacting with 3D virtual environments (Kerous and Liarokapis, 2016), but can also offer an opportunity to better understand and improve BCIs, to stimulate basic neurophysiological research, dedicated to investigation of the brain responses (Järvelä et al., 2015) and involved neural processes and behavioral research (Washburn, 2003). For example, BCI interconnected with virtual reality seems to be a promising approach to study specific brain responses, mental states and manipulating attention.

In this regard, another perspective application is in gamification of educational processes, which are currently undergo significant changes, leading to more enjoyable and effective personalized studying experiences.

### 1.3 BCI and VR

In the area of gaming recent research (Borhani et al., 2018; McMahon and Schukat, 2018; Cohen et al., 2016; Kerous and Liarokapis, 2016; Yan et al., 2016; Vourvopoulos and i Badia, 2016; Ferreira et al., 2014) as well as some past studies considered the possibilities of interaction of BCIs with virtual environments (Groenegress et al., 2010; Leeb et al., 2007; Lécuyer et al., 2008; Lotte and Guan, 2010; Leeb et al., 2007; Pfurtscheller et al., 2011). Most of these studies were again focused on applications for users with physical disabilities, whereas BCI-based interaction with virtual reality could also be an interesting experience for the user from the general public.

For example, gamification applied to education is able not only to increase engagement, but also motivation and performance (Hallifax et al., 2019).BCI in combination with AI is able to bring dynamic adaptation of educational tasks, depending on the personalized characteristics of the subject and subjects brain activity (Karkar, 2016; Zahabi and Razak, 2020; Abdessalem et al., 2018). This can allow educational system to keep balance between productivity and fatigue as well as between boredom due to the easiness of the task, engagement and stress (in case of hard tasking)(Dey et al., 2019).

At the same time, with an additional conjunction with VR it is promising to supply with proper surrounding, able to expand user experience: as for example in case of modelling complex processes, but also preventing distraction from outside processes, therefore holding attention.

### 1.4 BCIs and gamification

Since the first game, released to the market, there were many efforts applied to the use of BCIs in world-known games, such as “Tetris” (Pires et al., 2011), “Pacman”, “Pinball” (Tangermann et al., 2008), “World of Warcraft” (van de Laar et al., 2013), as well as many various home-made games utilizing various paradigms and nicely reviewed in (Kerous et al., 2018) and for consumer-grade EEG devices which are now becoming mainstream in (Vasiljevic and de Miranda, 2020).

Several BCI paradigms can be utilized in the area of gamification, regarding ways of controlling devices:

1. Neurofeedback (NFB), which exploit brain rhythms of different frequencies, such as theta (4–7 Hz), alpha (8–12 Hz), SMR (12–15 Hz) or beta (15–18 Hz) and challenge the user to control ongoing brain activity - typically the amplitude of particular brain rhythm or a different rhythm ratios, present in a visual, tactile or auditory feedback signals. This technology successfully used in treatment children with ADHD as it turned out that NFB has a better effect than traditional attention (Hurt et al., 2014) or working memory training (YuLeung To et al., 2016). Studies based on NFB training to improve cognitive skills most often involve SMR, beta1 and theta/beta ratio and show effectiveness after dozens of times of use (Egner and Gruzelier, 2004; Doppelmayr and Weber, 2011). Thus, it was showed that to consolidate developed skills, it is necessary at least 10 sessions or 3 hours of training (Gruzelier, 2014; Gruzelier et al., 2014).
2. Motor imagery (MI), which involves imagination of the movement of different parts of the body, resulting in sensorimotor cortex activity, detected as event-related EEG oscillations and transformed then into the signal sent to computer. BCI based on MI technology require repetitive training and can achieve quite high accuracy after several sessions (Prakaksita et al., 2016; Vasilyev et al., 2017; Škola et al., 2019). [(Vourvopoulos and i Badia, 2016)]
3. Or utilize different types of evoked potentials (EP), such as steady-state visually evoked potentials (SSVEPs) or P300 event-related potentials with different modalities. In contrast to rhythm-based neural interfaces, the BCI on the EP takes significantly less time and allows the user to manage commands from the first use (Kaplan et al., 2013). For example, SSVEP-based BCI can achieve accuracy above 95 % in just 5 seconds, which significantly exceeds the average accuracy of the system’s rhythms, obtained even after several training sessions (Erkan and Akbaba, 2018). Due to the simplicity of the occurrence of these potentials and the high accuracy of their detection, they are very easy to apply. However, SSVEP-based BCI cause a high degree of fatigue due to the constant high-frequency flashing of visual stimuli and do not lead to an improvement in attention.

These major paradigms can be employed alone or in various combinations, providing one an ability to create and implement active, reactive and passive BCIs (Zander and Kothe, 2011) based gamification of processes and routine operations.

### 1.5 P300 BCI

In application to games P300 is one of the most suitable paradigm, firstly used in speller application (Farwell and Donchin, 1988), nowadays also used for building the game plays (Congedo et al., 2011; Ganin et al., 2011; Edlinger and Guger, 2011).

P300 is an event-related potential (ERP) that occurs when a subject detects a significant stimulus in the context of the task (Picton, 1992). It’s called P300 because at the moment of recognition of the stimulus, a positive peak appears in the EEG in the region of 300 ms after its presentation. It’s also considered cognitive evoked potential because it’s caused not by the physical parameters of the stimulus, but by a person’s discrimination reaction to the relevant stimulus among others.

It has been shown that P300 BCI can be used by almost all healthy people, according to the study (Guger et al., 2009), 72.8% of participants were able to spell with P300 BCI with 100% accuracy, while only 3% were unable to spell correctly at all. P300 can also be successfully used by severely paralyzed patience without special training, which is usually needed for other paradigms such, as NFB or motor imagery, due to the fact that P300 resembles the natural functionality of the brain in goal selection and does not require acquaintance with something completely new (Farwell and Donchin, 1988).

P300 is widely used as a metric of cognitive functions by its presence, latency, amplitude and localization in both clinic and laboratory (Polich, 1986; 2007), for example, to measure attention for Parkinson disease and restless leg syndrome (Wang et al., 2000), Mild Cognitive Impairment and Alzheimer’s Disease (Papadaniil et al., 2016; Polich and Corey-Bloom, 2005), dementia (O’Donnell et al., 1992), mental workload (Causse et al., 2015).

In controversy to other deflections of evoked wave in response to target stimuli, such as N200, P300 mainly indexes the locus of attention but not eye gaze (Frenzel et al., 2011).

### 1.6 P300 BCI for cognitive training

The application of BCI, such as P300 in games or education not only leads to fun and new ways of interaction, increasing involvement and enjoyment, provided that BCI is easy to use, but can also substitute some of the communication mechanisms and lead to improvement of cognitive skills.

Being one of the fastest among currently available BCIs, and relating to a higher level attention process P300-based BCIs in games can not only be used for increasing levels of novelty and as a measure of mental workload, but also as cognitive training.

A study performed by (Rohani and Puthusserypady, 2015) demonstrated the use of P300 speller as attention training system, showing that P300 is connected to attention in the healthy subjects. In the other recent study P300-based speller BCI was used in neurofeedback training game, yielding in enhancement of ERP components, changing and increase in spatial attention tasks results (Amaral et al., 2015). There is one recent study of applying this type of interface as a tool for improving attention in healthy young adults (Arvaneh et al., 2019). In their work, authors modified standard P300-based speller BCI into a training game with individually increasing difficulty, prompting to generate stronger P300. It was shown that the training leads to an increase in the neural response to target stimuli, a greater suppression of the alpha-rhythm in experimental group and an improvement in the time of spatial attention task, following immediately after the training session.

These three studies were “proof of concept” preliminary studies, performed in laboratory conditions with wet electrodes, trivial gameplay and graphics as well as with lack of repetition nor investigation of the long-term effects.

In the present work we investigate an effect of P300 BCI-based VR-game with complex graphics on various cognitive functions of healthy adults, playing repetitive sessions with no-gel sponge electrodes and VR headset, using a panel of cognitive assessment tests.

## 2 EXPERIMENTAL DESIGN

### 2.1 Participants

In total, 53 healthy people were recruited for the study, however, some did not fully complete the training (all 5 sessions) and dropped out of the research. Therefore, the article presents data on 45 healthy subjects with normal or corrected-to-normal vision (25 females and 20 males) aged between 18 and 37 years old (mean age: 23,32 +-4,81 years) who participated in all stages of the experiment. All respondents were randomly assigned to three groups of 15 people each: experimental (P300+VR) (mean age 22,8 ± 5,24 years, 8 males), active control (VR game) (mean age 23,8 ± 4,38 years, 7 males) and passive control groups (VR movie) (mean age 24,0 ± 5,04 years, 8 males). Written informed consent was obtained from each participant after all the procedures were fully explained. Each subject was paid for participation in the study after completing all the stages of the experiment.

### 2.2 Experimental design (Procedure)

The overall experiment consisted of 5 sessions in a 2-week period and its structure can be found in the (Fig.1). Before the beginning of the experiment, all participants in all three groups performed a series of cognitive tests, which then was repeated after 1st, 3rd, and 5th training sessions. All of the participants answered state-assessment questionnaire, which has been presented before, in the middle and after each session.

**FIGURE 1.**
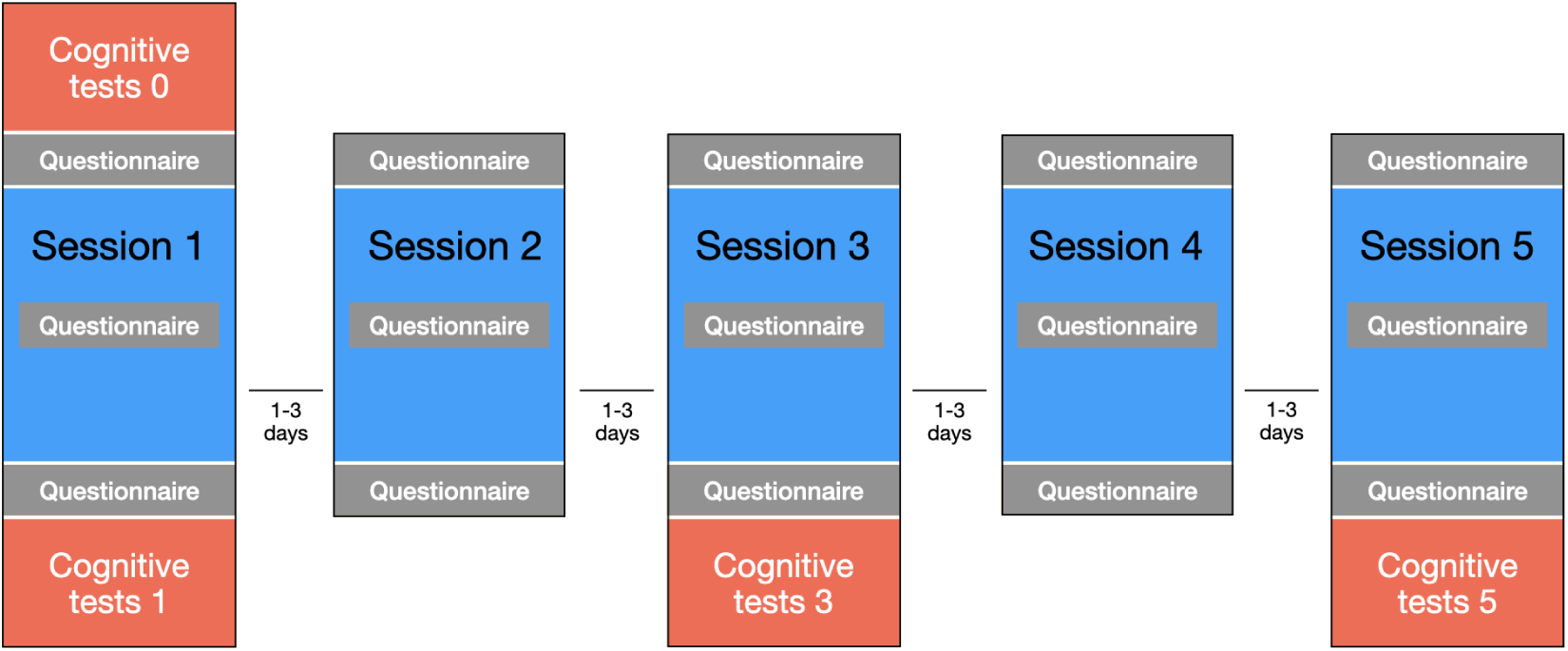
Experimental paradigm for all groups.

During each training session, participants performed their tasks described in 2.3 depending on the group. The duration of the session for each group was approximately 13 minutes. Virtual reality surrounding and stimuli were presented with HTC Vive Headset with two OLED panels for each eye, each with the resolution of 1080X1200, 110-degree field of view and a 90Hz refresh rate.

### 2.3 Tasks and Stimuli

Since the objective of the research was to study the effect of neurocontrol on cognitive functions, the main (principle) task of successfully passing the P300 game by experimental group participant was to focus his/her attention on the target stimulus while control groups were selected so that in one of them (an active control group playing VR game) respondents also needed to focus on the target stimulus but used traditional controller instead of BCI was involved, and task for another one - passive control group watching VR movie - was not based on utilizing attention at all.

#### P300 VR game

As a training task for the experimental group, a VR P300-based BCI game “Mind Fighters” (MF) (produced by LLC “Neiry”, Moscow, Russia) was chosen.

One MF game session consists of 6 similar rounds with 7 enemy robots appearing in front of the player’s avatar. Game structure is presented in the (Fig. 2). At the beginning of each session, a player passes short calibration procedure (Fig. 3 a). In each round it is necessary to make 5 shots. Before each shot, one robot flashes green once highlighting target stimulus (Fig.3 b). After this all 7 robots start flashing red in a random sequence but 7 times each. The player’s goal is to focus his/her attention on the target robot. The shot is made upon the target can be distinguished from non-target robots by classification by using visual ERPs including P300. Classification accuracy score for each shot is displayed as a feedback for player on the bar (Fig. 3 d). If the respondent’s concentration (attention on the target stimulus) was not well enough, a missfire occurs. If player’s attention was focused on non-target robot, then the shot is directed at this robot. EEG was recorded and accuracy was measured for each session of the game.

**FIGURE 2.**
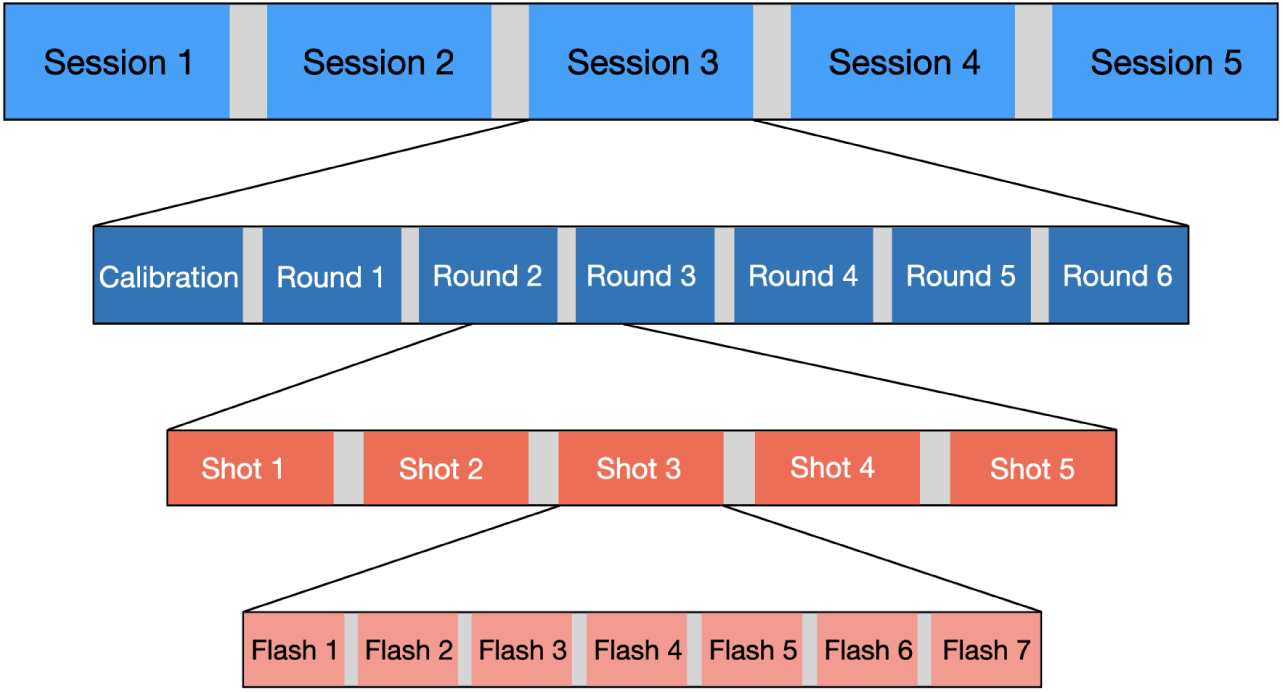
Experimental paradigm for P300 VR game.

**FIGURE 3.**
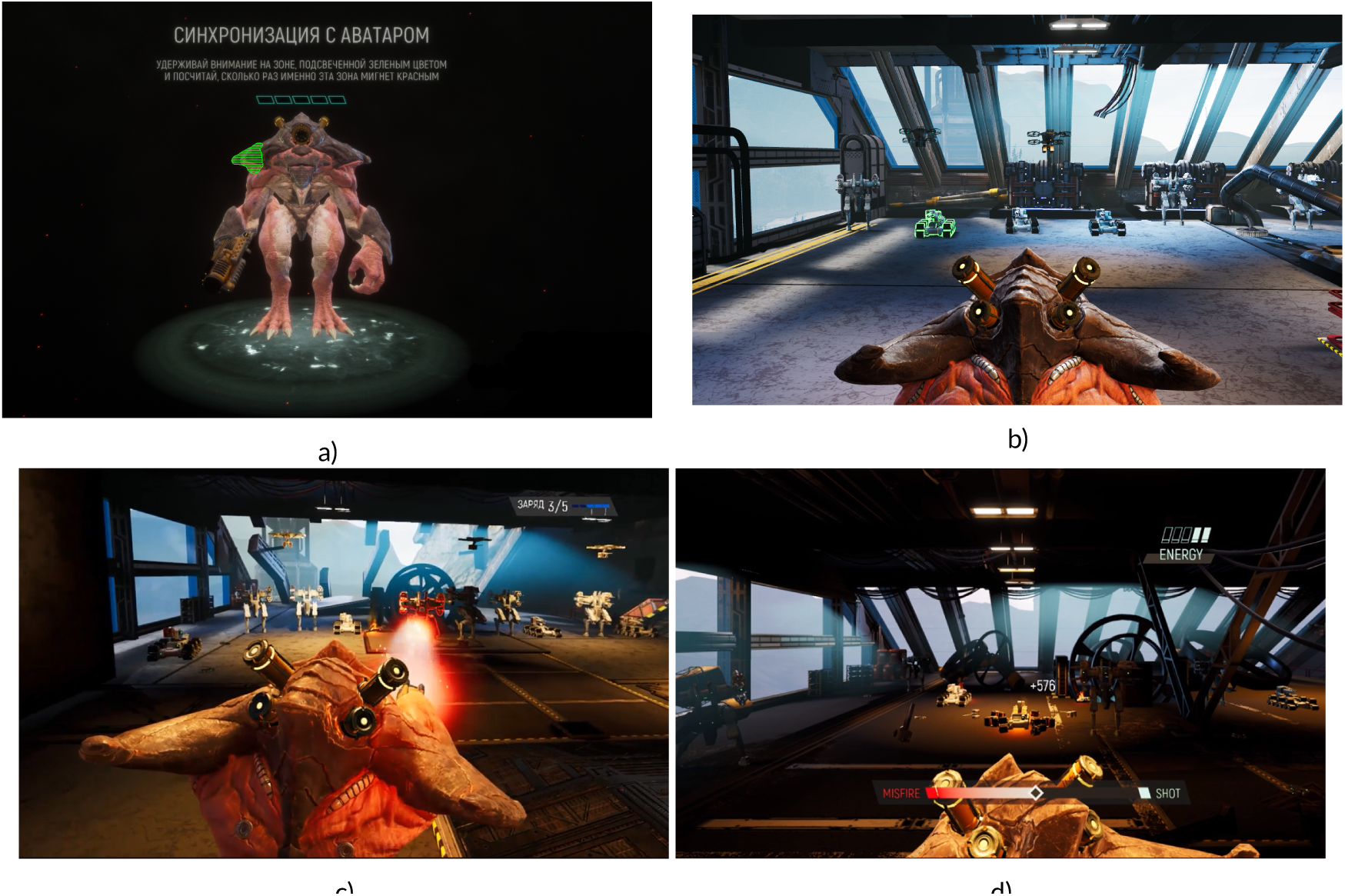
The gameplay of “Mind Fighters”. Calibration before the game (a), target activation (b), shot and estimation of the classification accuracy (c and d, correspondingly).

#### The active control group (VR game)

The active control group played “scientific” arcade VR game “InMind 2” where in contrast to experimental group game control is carried out by gaze direction detected by position of VR helmet (not eye-tracking) and not BCI. In order to select the desired object or to perform an action, the fixation of gaze for few seconds is necessary. However, the task of the game is similar to “Mind Fighters” : the player is located inside the avatar’s brain, and aims to collect target objects of correct color (yellow, green and blue brain cell bodies) by choosing them with gaze, at the same time avoiding non-target objects (red brain cells Fig. 4). Thus, here directing attention is again needed for successful completion of the game, although no specific concentration on the object.

**FIGURE 4.**
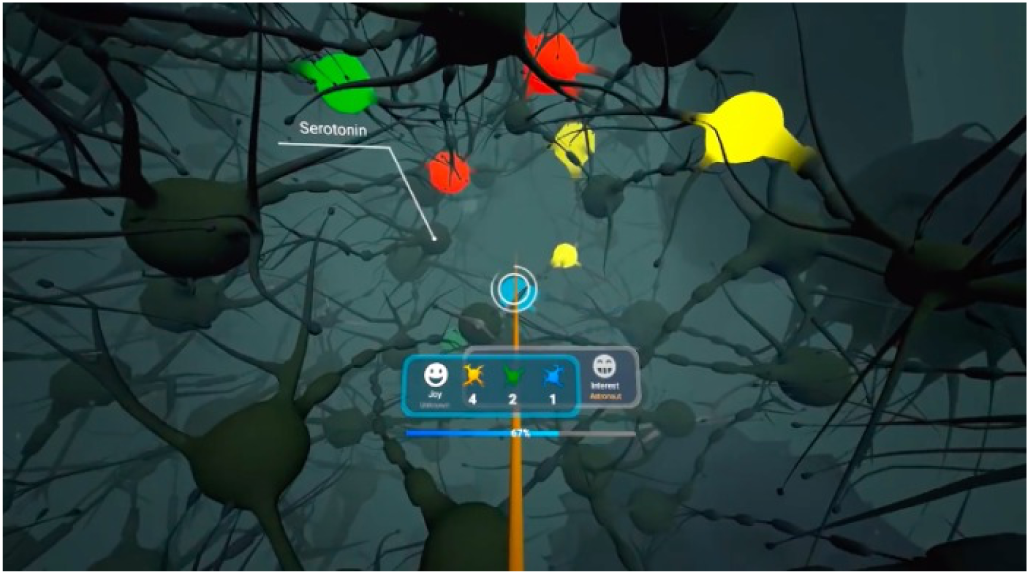
InMind 2 game representation. Yellow, green, blue and red-lighted targets (cell bodies) and a orange cone representing gaze direction for a choice.

#### Passive control (VR movie)

The passive control group watched a series of two fragments of VR movies without specific requirements of concentration or any other instructions, sitting in the chair.

### 2.4 Data Acquisition

#### EEG recording

EEG recording was performed using NVX-36 amplifier (Medical Computer Systems, Zelenograd, Russia) at 16 Ag/AgCl sintered electrodes placed according to 10-20 electrode system in frontal Fp1, central (C3, C1, Cz, C2, C4), centro-parietal (CP3, CP1, CP2, CP4), parietal (P1, Pz, P2) and occipital (O1, Oz, O2) lobes with the reference at A1 and ground at A2 (see Fig. 5) with 500 Hz sampling rate. Sponge-like material slowly releasing saline was used instead of conductive gel as the conductive material under the electrodes to provide a more user-friendly recording with a VR helmet.

**FIGURE 5.**
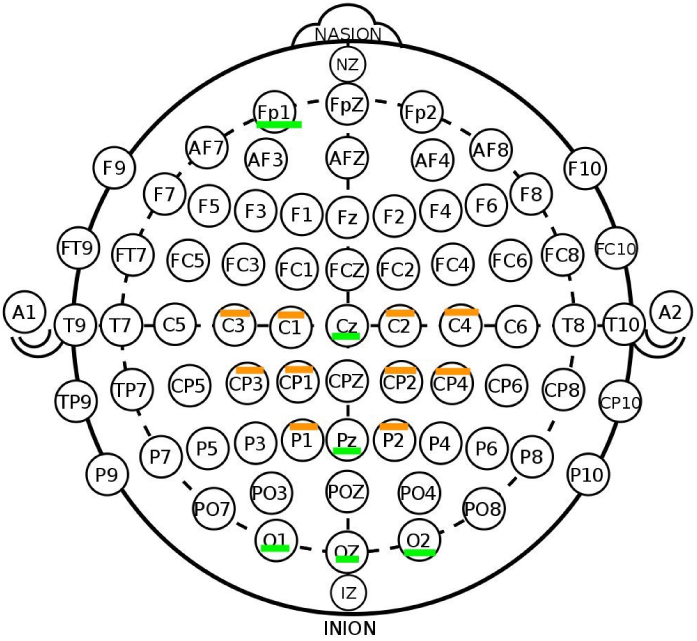
Configuration of electrode positions in the experiment.

EEG processing pipeline line consisted of butterworth digital filtering of 4th order with cutoff frequencies of 15 and 0.5 Hz. Filtering was applied in a forward-backward manner. After that, a 10 times decimation was performed. Scale EEG values were also clipped to 150 micorvolts. On the next step, CSP were used to amplify signal and reduce artifacts impact. Classification was performed with LDA classifier shrinking data using the Ledoit-Wolf lemma.

#### Cognitive assessment

A panel of tasks was prepared for the experiment for cognitive function evaluation. All subjects performed adapted Eriksen Flanker tasks, Visual search task, Go/no-go task (with visual stimuli) and Corsi block tapping task on the computer. Reaction times and error rates were recorded with the presentation software. Behavioral data were then analyzed and compared between the groups with the use of statistical tests.

**TABLE 1.** The list of cognitive assessment tests and measured parameters

| test | Cognitive function | parameter measures |
| --- | --- | --- |
| Eriksen Flanker test* | Attentional inhibition | time, accuracy |
| Visual search** | Visual spatial attention | time, accuracy |
| Go/no-go task | Response inhibition | time, accuracy |
| Corsi block tapping task | Working memory | num. of remembered objects |

\* - congruent and incongruent; \*\* - 5, 10, 15 and 20 elements as distractors.

**Eriksen Flanker task** is designed to study the cognitive processes of stimulus detection and recognition under conditions of distracting information or “noise” (Eriksen and Eriksen, 1974). Flanker task is used as a measures of selective attention when it is necessary to evaluate top-down regulation of attention (Posner and DiGirolamo, 1998; Fan et al., 2003; Rueda et al., 2005). It allows to estimate the ability to inhibit reactions that are inappropriate for a particular context. Thus, the Flanker test is a measure of attentional inhibition. Two types of stimuli are presented in the test: task-relevant (target) and task-irrelevant stimuli (non-target). Combination of correspond task-relevant and task-irrelevant stimuli represents congruent trails, whereas combination of unrelated stimuli becomes incongruent trails.

**Visual search** is a type of perceptual task requiring attention that typically involves an active scan of the visual environment for a particular object or feature (the target) among other objects or features (the distractors) (Treisman, 1977; Treisman and Gelade, 1980). Visual search relies primarily on endogenous orienting because participants have the goal to detect the presence or absence of a specific target object in an array of other distracting objects.

**Go/No-go** paradigm was developed to test the ability to perform an appropriate reaction under time pressure and to simultaneously inhibit an inappropriate behavioural response (Criaud and Boulinguez, 2013; Verbruggen and Logan, 2008). In this form of behavioural control it is important to suppress a reaction triggered by an external stimulus to the benefit of an internally controlled behavioural response. In this paradigm, the focus of attention is directed to predictably occurring stimuli that require a selective reaction, that is, to react or not to react.

**The Corsi block-tapping test** allows to assess visuospatial short term working memory (Corsi, 1973; Kessels et al., 2000). It involves mimicking a researcher as they tap a sequence of up to nine identical spatially separated blocks. The sequence starts out simple, usually using two blocks, but becomes more complex until the subject’s performance suffers.

## 3 RESULTS

### 3.1 Playing P300 BCI Game

#### Neurogaming and performance

During game sessions most of participants were able to successfully use P300 BCI controller for playing “Mind Fighters”. All the 15 subjects were able to play at least one game round with 100 % accuracy, however with different average results in overall performance. The online overall accuracy varied from 30 to 100 % in one game session (consisted from 6 rounds) with the mean accuracy across all the subjects and all the sessions at the level of 69.3 ± 18.7 % (averaged accuracy for each session are presented in the Fig. 6). The best performance was found from subject 12 with an error rate of 0.062 across all the 30 trials in 6 rounds game play over 5 days. (starting from 0.766 accuracy in 1st training session up to 1, 0.966, 1, 0.933, 0.966). The worst result was showed by participants 4 and 1, who performed with 0.34 and 0.393 accuracy correspondingly. Classification accuracies table is present in the Fig.17.

**FIGURE 6.**
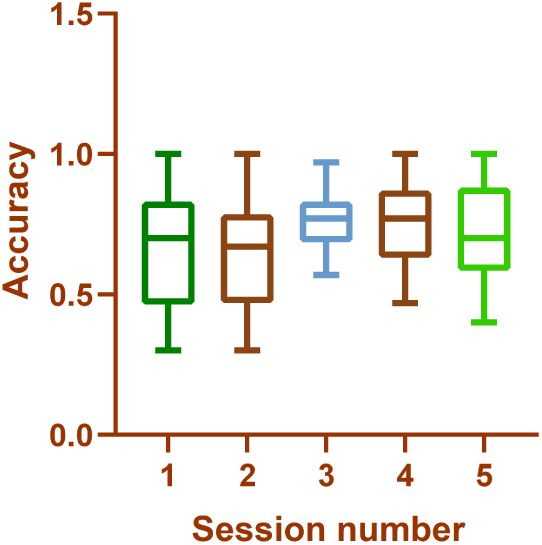
Boxplots of classification accuracy in each training session. Whiskers represent min and max value.

Self-estimation of psychological state showed that on average playing the game led to slight increase of mood, concentration, however also level of general tiredness and eye strain. The same tendencies were observed in results of self-estimation for active and passive control groups. Overall, playing the neurogame at the same time was perceived as an interesting experience and led to positive feedback from participants.

### 3.2 ERPs

Analysis of the recorded EEG revealed the presence of P300 wave at all the electrodes’ locations in frontal, parietal, central and occipital areas. Signal at some characteristic for P300 potential locations (Cz and Pz) is displayed in the Fig. 7. With a distinct increase of the amplitude of the wave in response to target stimuli in comparison to nontarget activation, it indicates correct P300 based distinguishing between target and non-target activations used for classification in the “Mind Fighters” game. Some individual differences in the evoked potential across subjects can be clearly seen in the images (see Fig. 8). Subjects 2, 6 and 8 were picked up for illustration out of all participants because they nicely represent different shapes of P300 potential.

**FIGURE 7.**
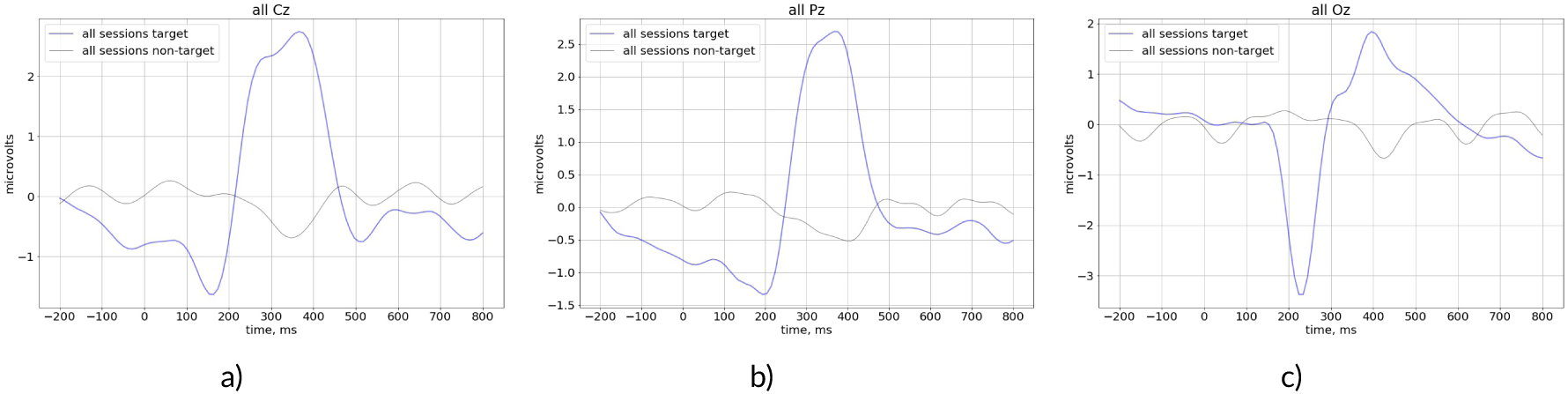
Grand averaging of EEG signal at a) cz, b) pz c) oz locations.

**FIGURE 8.**
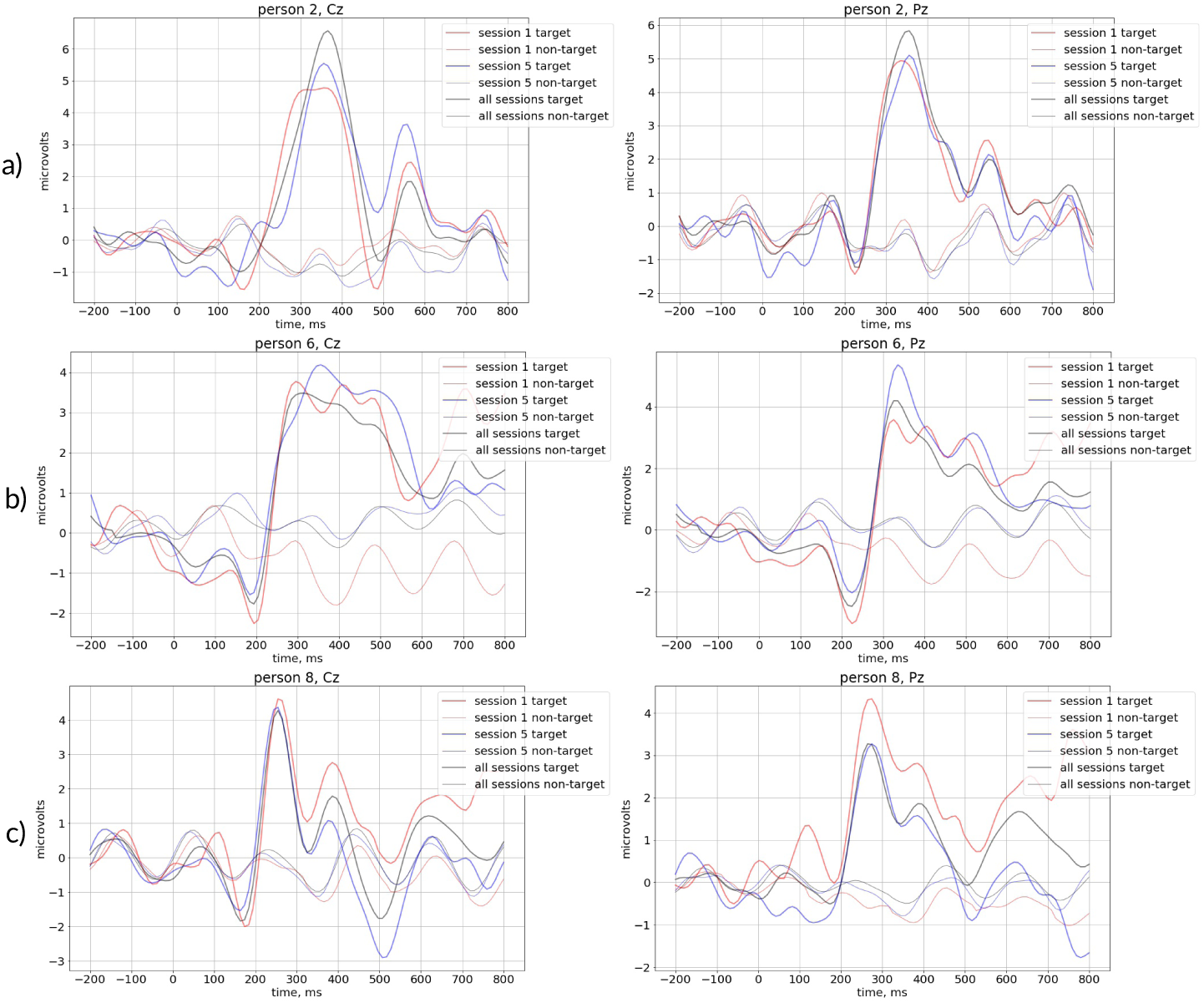
P300 at main locations: comparison of target-non target cz and pz.

These variations of amplitude and latency as well as a ERP wave shapes are one of the reason of complication of classification process, decreasing the accuracy. We also observed a high amplitude negative wave at occipital location in response to target stimuli (Fig. 7 c and Fig. 9). It was found out, that latency and amplitude of P300 were stable for the most of the participants, however in some subjects we have detected increase of P300 amplitude from 1st to 5th training session with simultaneous decrease in standard deviation of signal amplitudes among particular trials (see Fig. 10).

**FIGURE 9.**
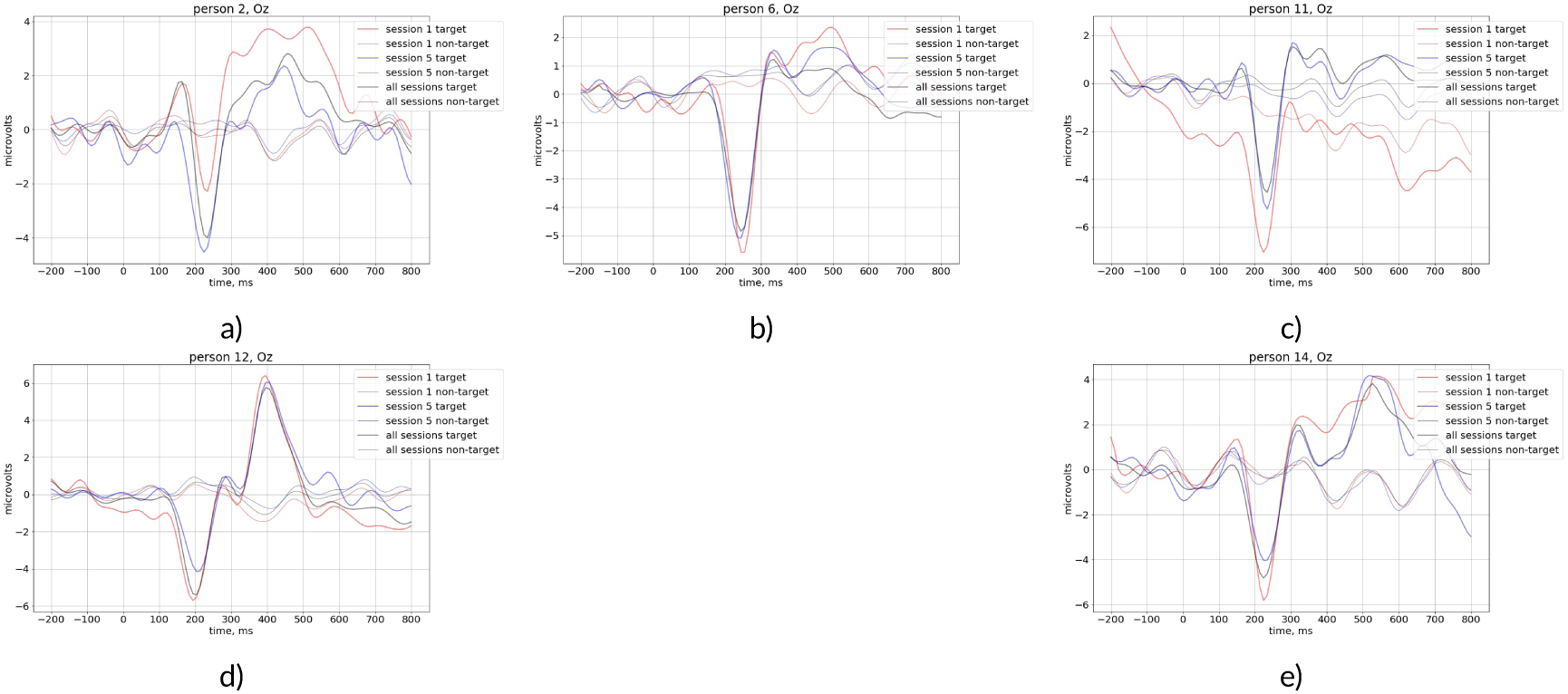
A typical for P300 BCI negative peak appearing in response to target stimuli at the occipital location detected in EEG of different subjects.

**FIGURE 10.**
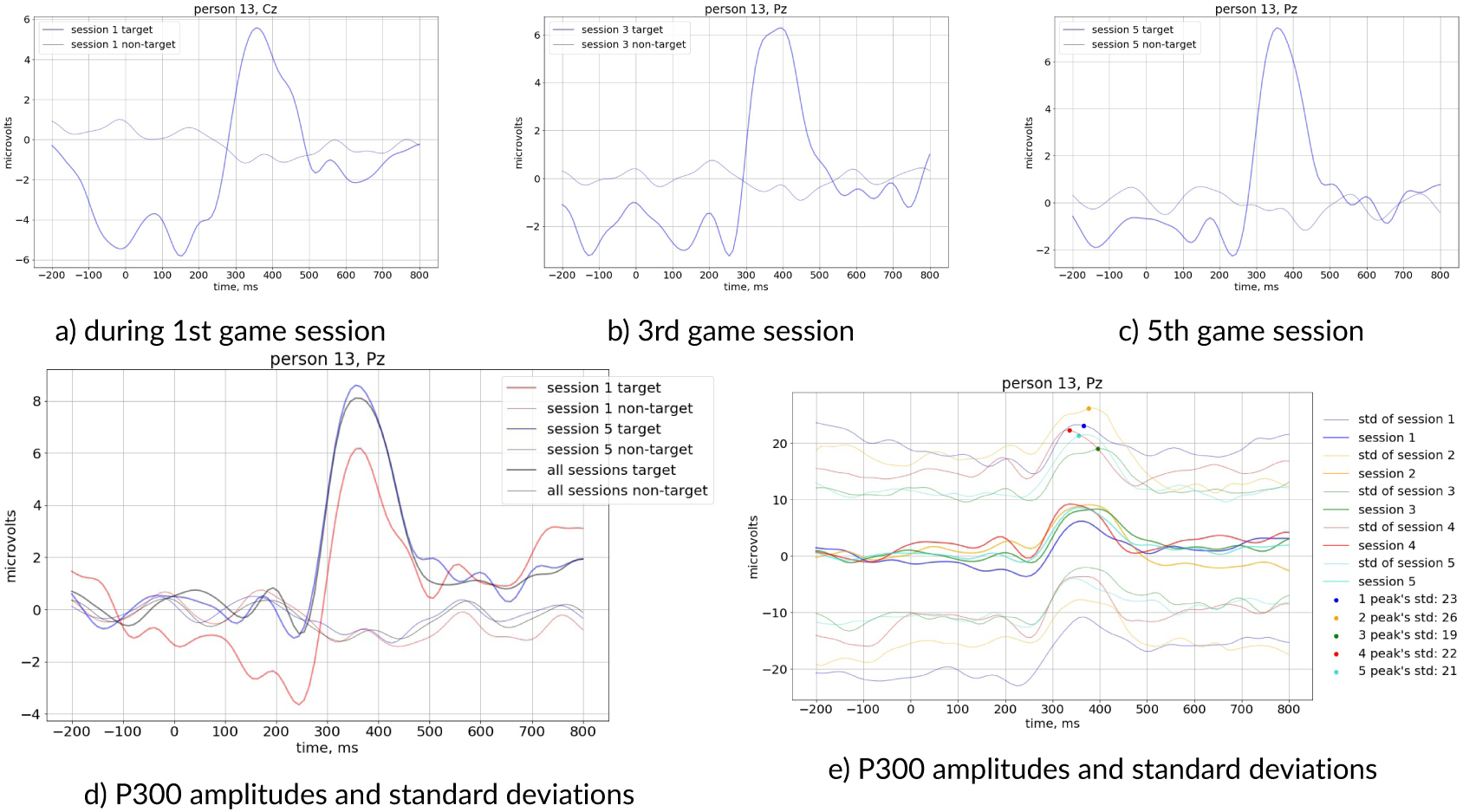
Averaged P300 amplitudes during a) 1st session 3rd session, c) in 5th session increase in P300 amplitude and decrease in standard deviations e).

### 3.3 Cognitive assessment

Results of the cognitive assessment reveal changes in cognitive tests performance between experimental and both control groups. Over all measured characteristics, those presented in the Fig. 11, showed the most change of cognitive functions in subjects from experimental group (detailed results with extensive data are presented in Fig. 16.)

**FIGURE 11.**
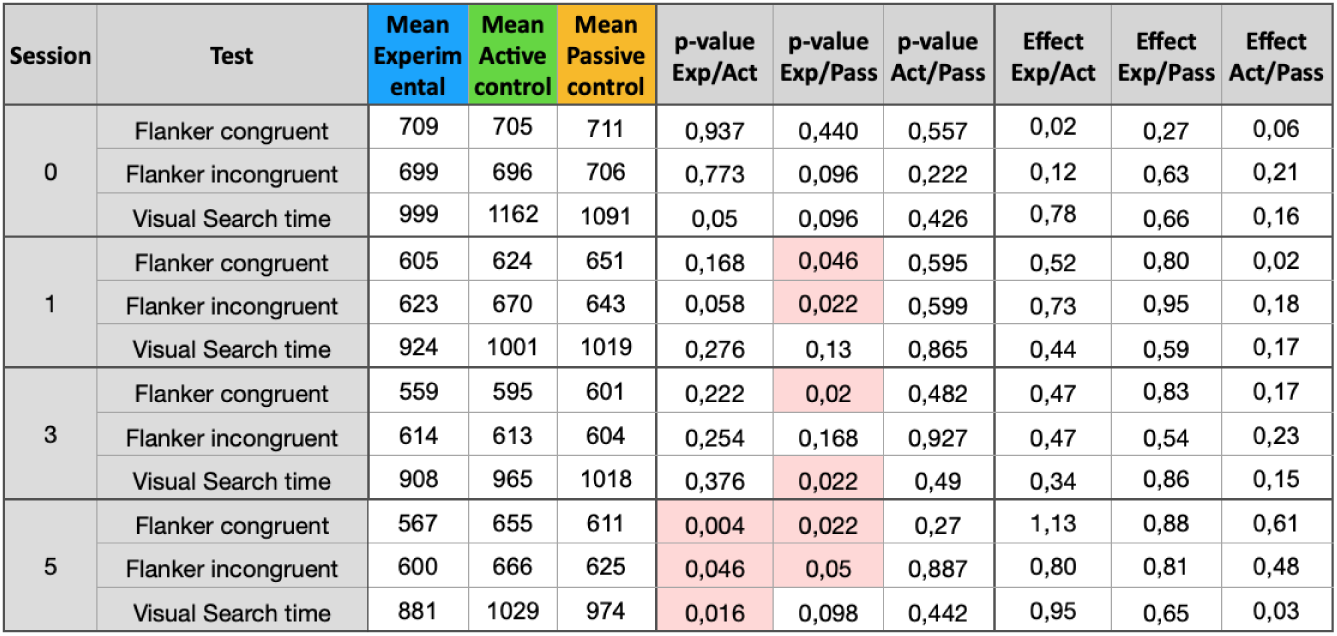
Cognitive assessment differences between experimental and control groups. Reaction time (prior 1st session, after the first, third and fifth sessions - 0, 1, 3, 5, respectively), test, average values for experimental, active and passive control groups, p-values (permutation test) and size effects (Cohen’s d).

**FIGURE 12.**
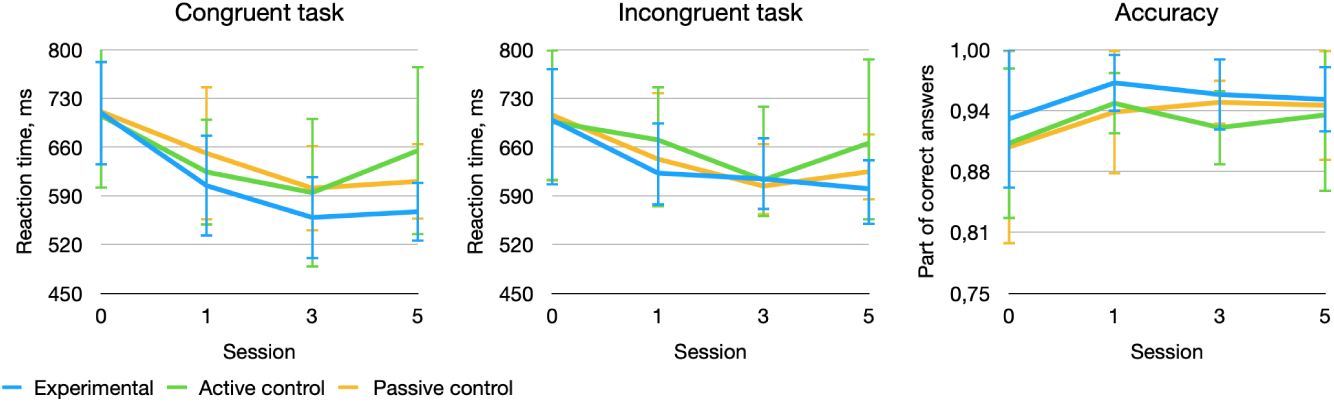
Cognitive assessment differences between experimental and control groups. Average reaction times in flanker task (congruent and incongruent conditions) and average accuracy of the response prior 1st, after the first, third and fifth sessions - 0, 1, 3, 5, respectively. Error bars show standard deviation.

**FIGURE 13.**
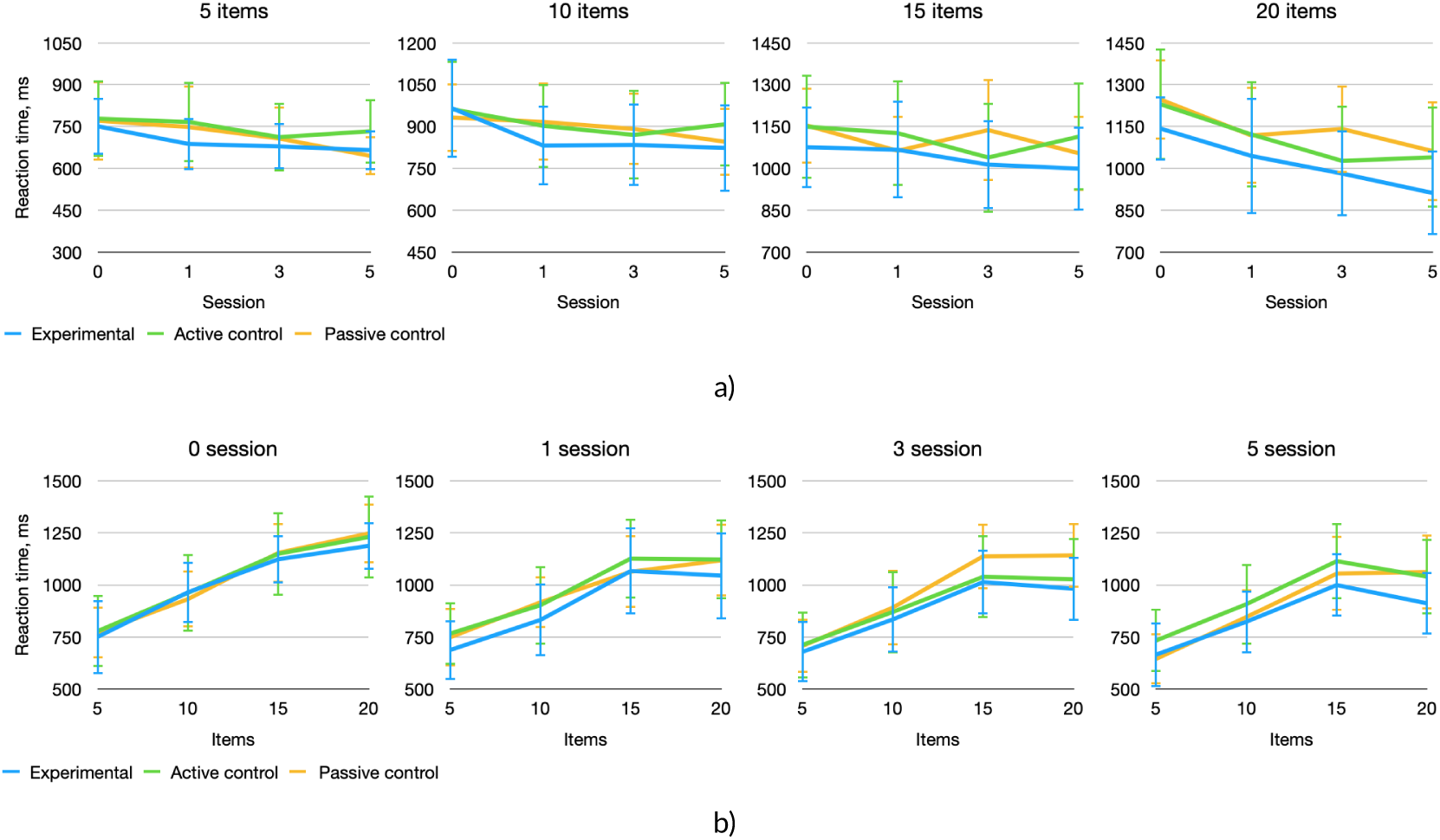
Cognitive assessment differences between experimental and control groups. Average reaction times in visual search task depending on session number for 5, 10, 15 and 20 items presented in the test (top line); and average reaction times depending on the number of distractors at baseline (0), after 1, 3 and 5 sessions of P300 BCI training (bottom line). Error bars show standard deviation.

**FIGURE 14.**
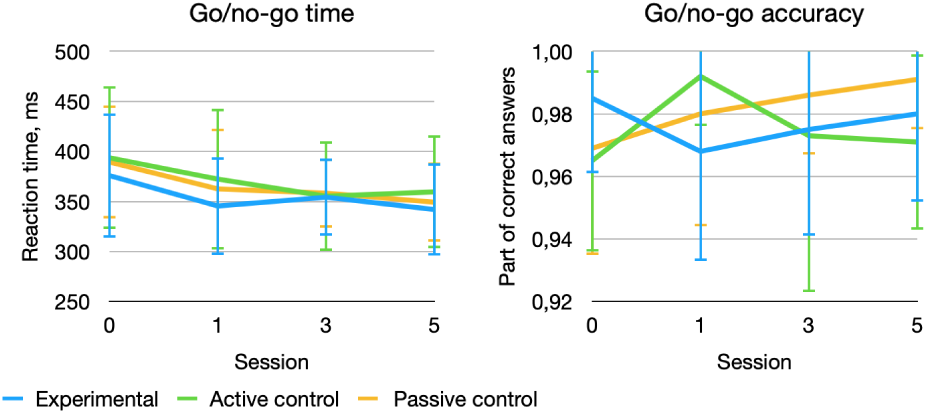
Cognitive assessment differences between experimental and control groups. Average reaction times and accuracy in Go/no-go task at baseline (0), after 1, 3 and 5 sessions of P300 BCI training. Error bars show standard deviation

**FIGURE 15.**
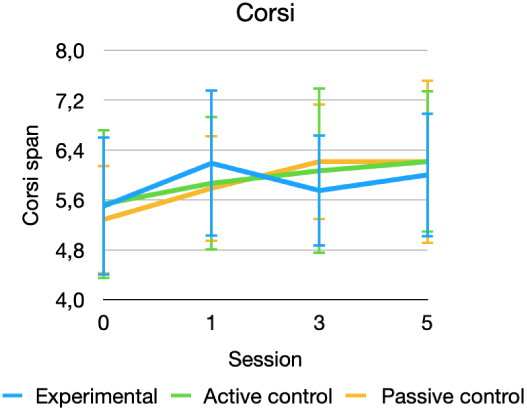
Cognitive assessment differences between experimental and control groups. Mean scores of the Corsi Block Test for the experimental and control groups at baseline (0), after 1, 3 and 5 sessions of P300 BCI training. Error flags indicate standard deviations.

**FIGURE 16.**
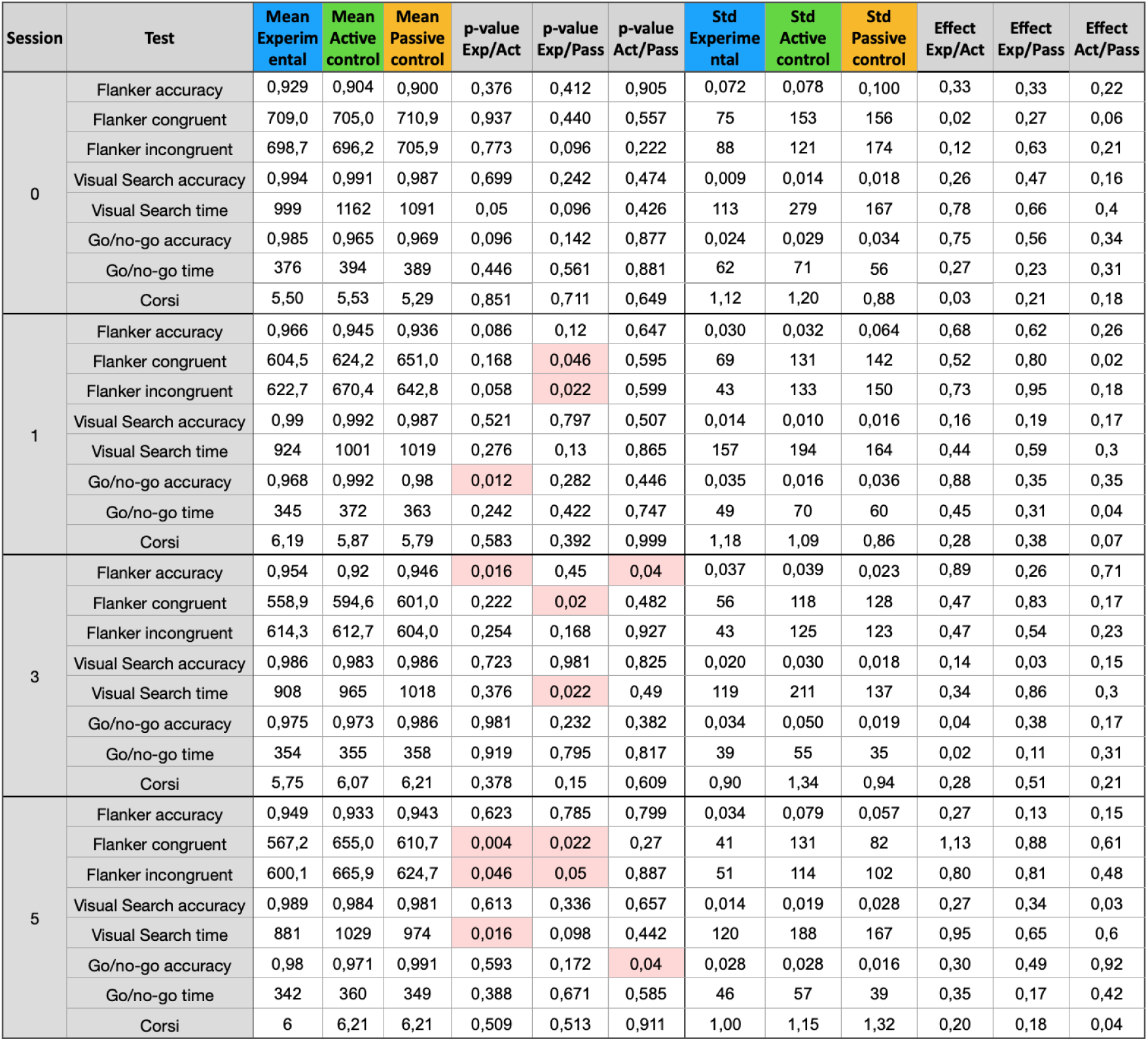
Cognitive assessment results for experimental and control groups with mean values, standard deviations, calculated p-values and Cohen effects

**FIGURE 17.**
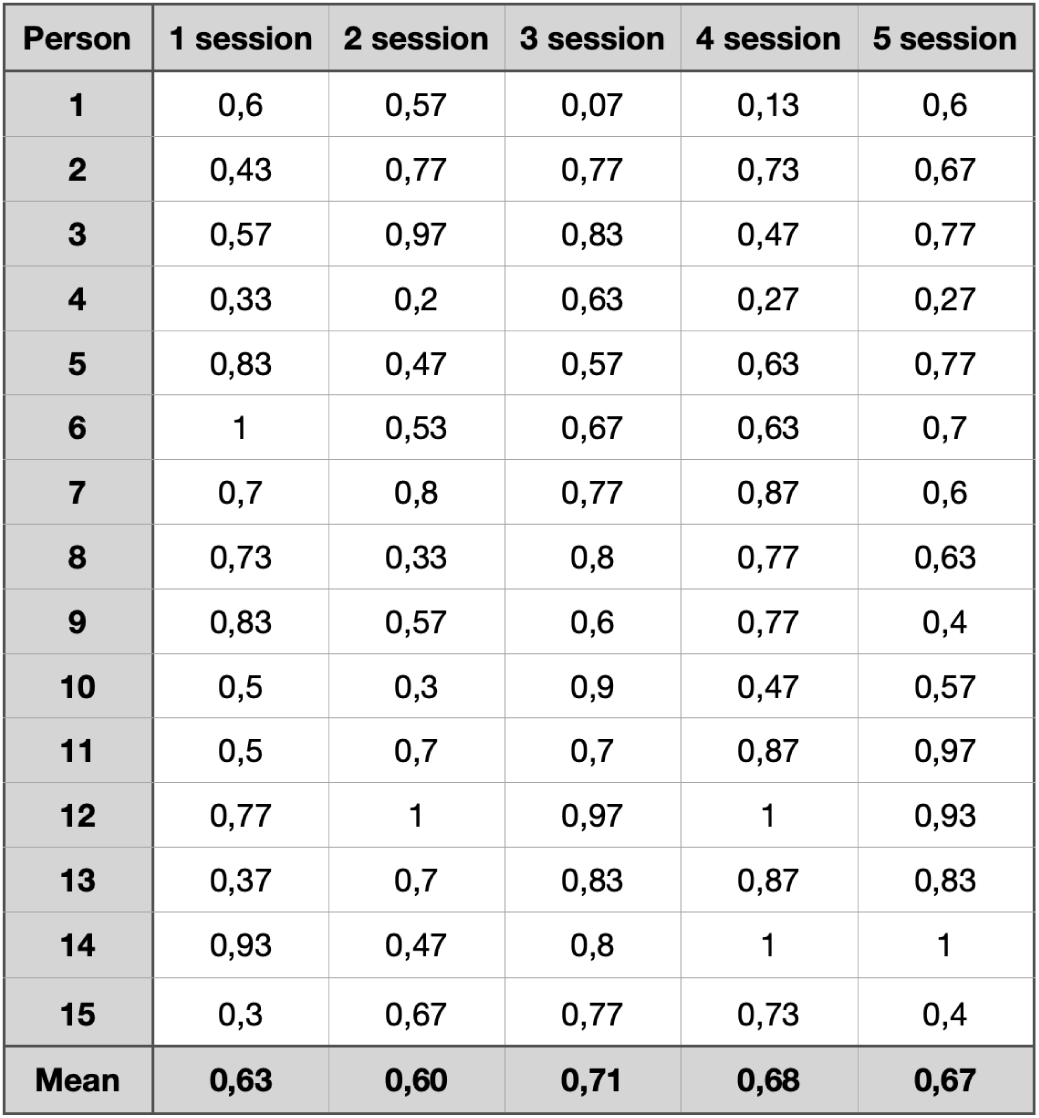
P300 game accuracy for experimental group participants. Percentages for each session is the average over 6 game rounds.

It was found out that attention assessment in reaction times has undergo the biggest change, while other parameters including accuracies and working memory tests on average didn’t show much change in respondents.

#### Eriksen Flanker task

One of the tasks with distinct changes between experimental and control groups appeared to be Eriksen Flanker Task, which consisted of congruent and non-congruent subtasks, related to distinguishing a target stimulus in the surrounding of items of the same or opposite groups. Results of reaction times and accuracies for this tasks are presented in the Fig. 12. While no significant difference was observed between experimental and control groups in the beginning of the experiment (at the baseline), the 5th session revealed statistically significant differences in task performance (p-value is present in the table), indicating more progressive learning by experimental group.

A typical difference in reaction times was also observed between congruent and non-congruent condition, where results of incongruent tasks are higher (see Fig. 12 a. and 12 b.) due to the flanker effect (Eriksen and Eriksen, 1974).

It was also revealed that improvement in the time taken for tests by the experimental group was not accompanied by a drop in the accuracy of the answers (Fig. 12 c). Moreover, the accuracy of the experimental group was slightly higher than that of the control groups.

**TABLE 2.** Effects of Mixed ANOVA results for Eriksen Flanker task

| Task | Group | Session | Interaction |
| --- | --- | --- | --- |
| Congruent | $F = 1.40, p = 0.259$ | $F = 28.49, p = 0.000$ | $F = 1.95, p = 0.079$ |
| Non-congruent | $F = 1.37, p = 0.268$ | $F = 13.61, p = 0.000$ | $F = 1.31, p = 0.261$ |

#### Visual search

Improvements in performance in Visual Search task are found to be in overall consistency with Flanker task results. A similar decrease in time from 0 to 5th session is clearly observed, indicating the effect of learning (Fig 13 a)).

The Visual search test in the current study included several difficulty levels presented randomly during the task. The complication was in an increase of simultaneously presented stimuli, among which it was necessary to find the target. Therefore the total number of stimuli that appeared at one time on the screen was 5, 10, 15, or 20. The increase in irrelevant objects in the search task complicates the process of scanning space and thereby creates a large load for spatial attention. Considering the uneven distribution of attention and time for solving a task with a different number of elements, the data of the results of the test time were distributed and analyzed according to the number of stimuli presented: 5, 10, 15, 20 items. (see Fig. 13 a)).

A more detailed examination revealed tendencies for a more significant reduction in the time taken to solve the complicated task of visual search in the experimental group (Fig.13). Based on this trend, it can be assumed that training on the P300 can improve spatial attention and orientation in a very noisy environment, however, this assumption needs to be verified on a larger sample of respondents.

**TABLE 3.** Effects of Mixed ANOVA results for Visual search task

| Items | Group | Session | Interaction |
| --- | --- | --- | --- |
| 5 | F = 1,21, p = 0.309 | F = 4.65, p = 0.004 | F = 0.67, p = 0.675 |
| 10 | F = 1.29, p = 0.286 | F = 10.51, p = 0.000 | F = 1.07, p = 0.385 |
| 15 | F = 2.52, p = 0.093 | F = 7.55, p = 0.000 | F = 1.88, p = 0.089 |
| 20 | F = 2.86, p = 0.068 | F = 12,83, p = .000 | F = 2.42, p = 0.030 |

**TABLE 4.**

| Abbreviation | meaning |
| --- | --- |
| Flanker congruent | time of congruent answers of the Eriksen Flanker task |
| Flanker incongruent | time of incongruent answers of the Eriksen Flanker task |
| Flanker accuracy | accuracy of the Eriksen Flanker task |
| Visual search time | reaction time of the Visual search |
| Visual search accuracy | accuracy of the Visual search |
| Go/no-go time | reaction time of the Go/no-go task |
| Go/no-go accuracy | accuracy of the Go/no-go task |
| Corsi | the maximum number of correct answers in Corsi block tapping task |
| Mean | the average value of the index for the group |
| p-value | significance level |
| Effect | Cohen's effect size |
| Std | standard deviation |
| Exp | experimental group |
| Act | active control group |
| Pass | passive control group |

Legend for the table in Fig. 11 and Fig. 16 .

#### Go/no-go task

Statistical analysis of the results did not reveal significant differences in terms of time and accuracy of passing the Go/no-go task. All the participants regardless of the group performed the task with high accuracy and similar response times. It should be noted that both indices of this test showed a ceiling effect for all three groups and therefore we can conclude that these tests appeared to be too simple for evaluation of healthy adult respondents. Thus, differences in the cognitive indicator of impulsivity estimated by go/no-go were not identified.

#### Corsi block tapping task

According to the results of the statistical analysis of Corsi block tapping task performance, no significant differences were found between the experimental and control groups of respondents. It is likely that it is more appropriate to change methodology to assess visuospatial working memory: increase difficulty or apply more advanced tests, which could give more informative and variable data. Thus, according to the results of this test, no effect of training on working memory was shown.

## 4 DISCUSSION

The data we obtained during the experiment is consistent with previous studies. The majority of participants were able to utilize P300 BCI, with a rather high-performance (Guger et al., 2009). Similarly to other studies (Shishkin et al., 2009) it was found out the presence of specific for P300 BCI high-amplitude negative peak (N1), located at occipital electrodes approximately 200 ms after target stimuli. Comparison of the characteristics of P300-wave (latency and amplitude) over 5 sessions demonstrates slight adaptation from the first game recordings (Ganin et al., 2013) and overall stability of characteristic of this ERP component. A slight increase of P300 amplitude with a simultaneous decrease of standard deviation from 1st to 5th session, detected in some participants could be associated with the improvement in the evocation of ERPs in these healthy subjects, induced by training and increase in attention (Rohani and Puthusserypady, 2015). However, in most of the participants, the P300 amplitude and latencies were stable.

The overall tendency to a general decrease of reaction time from session to session observed in all the cognitive tests used in the experiment (Fig. 12, 13,14), is explained by the respondents’ learning to pass the test. However, it should be noted that the time taken for these tests by the experimental group is stably smaller than that in the control groups. Moreover, a facilitated improvement is observed for the experimental group as evidenced by the initial angle of the curve.

The most important result is a significant difference between the experimental group and both control groups with no significant differences between control groups after the 5th session Eriksen Flanker task, used to measure attentional inhibition as an ability to resist interference from distracting stimuli (Nigg, 2000; Friedman and Miyake, 2004; Wiebe et al., 2008; Kane et al., 2016; Tiego et al., 2018). Therefore our data allow us to assume that the respondents from the experimental group showed significant improvements in attentional inhibition (concentration), associated with performance in Flanker task in comparison to control groups.

Additionally, some differences were found in another attentional task - visual search task, dedicated to the evaluation of spatial attention. Besides these results with the most significant differences in a congruent flanker task, no other test from our cognitive panel did reveal significant changes. The evaluation methodology was chosen based on the assumption that testing would continue at different age groups and it turned out that test assignments were too easy for adult participants.

Taking into account the similarity of tasks for the experimental and active control group as well as similar surroundings, which was carefully chosen for the experiment, it can be noticed that significant differences in Eriksen task performance between experimental and both control groups after 5 training sessions appear due to 2 factors: natural learning and impact of the use of P300 BCI.

Results of Mixed ANOVA for both attentional tasks showed that the main factor influencing the improvement of the attentional tests’ reaction times is ‘session’. This can be explained with participants’ ability to learn how to solve cognitive tasks and therefore is associated with implicit learning. However, differences in results between experimental and two control groups after 5th gaming sessions together with statistical analyses suggest the increase in the impact of P300-training on changes in cognitive assessment from session to session.

Therefore obtained data suggest a more obvious effect after increase the number of training sessions when adaptation to a task is completed and other factors start playing a more significant role.

Given the p-value, the size effect, and standard deviations for evaluated parameters (Table 11), we can assume that with the increase in participant number, a more strong correlation can be observed. Some previous research together with basic knowledge supports our suggestions. The similarity of a Flanker task and a P300 BCI paradigm in the involvement of attentional inhibition either to recognize stimulus in the environment of similar stimuli (Eriksen and Eriksen, 1974; Tipper, 1985; Friedman and Miyake, 2004; Nigg, 2017) or to select one target stimulus with focusing attention, can link p300 BCI, also involving memory (Polich, 2007), with better performance in attentional tasks.

## 5 CONCLUSION

This study investigated whether P300 BCI gaming can influence cognitive functions of healthy adults and evaluated the experience of playing BCI games in a rich VR environment without using wet electrodes. We have found that participants have an interest in the BCI-VR system and enjoyed gaming. Significant changes in cognitive assessments were shown after 5 experimental sessions for the experimental group in comparison to both control groups in the tasks, associated with inhibition and spatial attention. It was found that natural learning has more impact on the results of cognitive tests, however, results of the study suggest that effect of P300 BCI is present and can become more visible with the extension of training timescales and increase in group sizes, aiming to countervailing strong effects of individual differences and task adaptation. Results of our preliminary study suggest that P300 training has a positive effect on selective attention and the ability to inhibit distracting stimuli and quantitative evaluations of BCI impact need to be estimated in further research.

## Acknowledgements

Authors would like to thank scientific consultants: Sergey Shishkin, Marie Arsalidou and Alexey Kotov who helped with experimental design and/or discussion, as well as Alexey Khalezov for helping with management during all the stages of the experiment.

## Conflict of interest

MB declares no conflict of interest. AK and AS are employees at Neiry LLC. AP is Chief Executive Officer at Neiry.

**A. One** Please check with the journal’s author guidelines whether author biographies are required. They are usually only included for review-type articles, and typically require photos and brief biographies (up to 75 words) for each author.

## Notes

### Competing Interest Statement

declared in the manuscript

### Summary of Updates

revision of several sections and language improvements

